## Supplementary Information for "Multiple blood feeding in mosquitoes shortens the *Plasmodium falciparum* incubation period and increases malaria transmission potential"

### SI Text

#### Modeling Details

Our simplified formula for $R_0$,

$$R_0(T) = \sqrt{f(T)e^{-g(T)}},$$

can be related to the original equation from Mordecai *et al.* (1) reproduced here as,

$$R_0(T) = \sqrt{\frac{a(T)^2 bc(T) e^{-\frac{\mu(T)}{PDR(T)}} EFD(T) p_{EA}(T) MDR(T)}{Nr\mu^3(T)}},$$

by

$$f(T) = \frac{a(T)^2 bc(T) EFD(T) p_{EA}(T) MDR(T)}{Nr\mu^3(T)}$$

and

$$g(T) = \frac{\mu(T)}{PDR(T)}.$$

The expression is based on temperature-dependent trait data from *Anopheles* species that vary with temperature including biting rate (*a*), vector competence (*bc*), adult mosquito mortality ($\mu$), parasite development rate (*PDR*), egg-to-adult survival probability ($p_{EA}$), mosquito development rate (*MDR*) and eggs laid per female per day (*EFD*). Note that adult mortality is calculated through daily adult survival (*p*) via $p = e^{-\mu}$; parasite development rate is one over the extrinsic incubation period (*EIP*); and mosquito development rate is one over the larval development time ($\tau_{EA}$). Human related quantities—human density (*N*) and recovery rate (*r*)—are not temperature-dependent. Furthermore, explicit values are not necessary as *N* and *r* cancel out in our final ratios. The temperature-dependent traits of biting rate, parasite development rate and mosquito development time were fit to Briere functions, represented by

$$cT(T - T_0)(T_m - T)^{\frac{1}{2}},$$

where *c*, $T_0$ and $T_m$ are constants defined in previously published work and replicated here for completeness (**Table S5**) (1). The temperature-dependent traits vector competence, daily adult survival, egg-to-adult survival probability and eggs laid per female were fit to quadratic functions, represented by

$$qT^2 + rT + s,$$

where *q*, *r*, and *s* are constants determined in previously published work and replicated here for completeness (**Table S6**) (1).

Our modified basic reproductive number incorporating a second blood feeding by scaling EIP using the term $\beta$, which we refer to as $R_0^b$ is given by

$$R_0^b(T) = \sqrt{\frac{a(T)^2 bc(T) e^{-\beta\frac{\mu(T)}{PDR(T)}} EFD(T) p_{EA}(T) MDR(T)}{Nr\mu^3(T)}}.$$

We determine the scaling parameter $\beta$ as the relative reduction in EIP in the presence of a second blood feed, given by

$$\beta = \frac{2\ blood\ feeds}{1\ blood\ feed} = \frac{8.63}{10.88} = 0.73.$$

The change in $R_0$ using a shortened EIP is shown by the ratio of the modified $R_0$ to the original $R_0$ as

$$\frac{R_0^b}{R_0}.$$

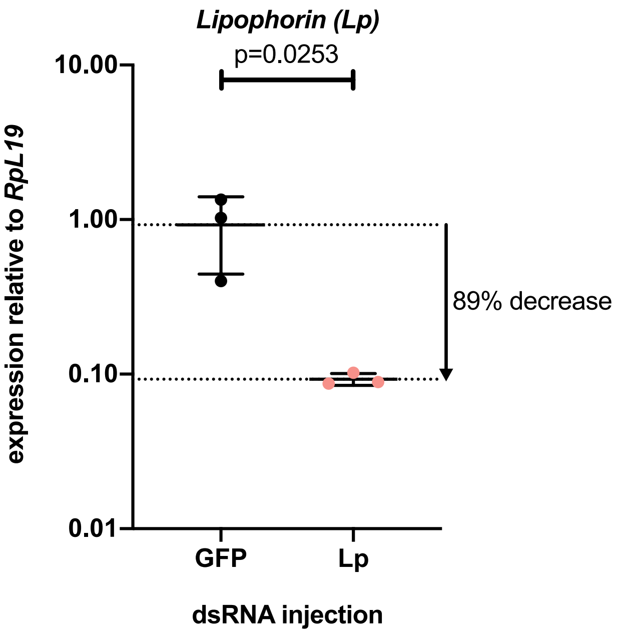


**Fig. S1. *Lp* expression is effectively silenced following ds*RNA* injection.** *Lp* gene expression was determined in pools of 5-10 decapitated females at 6 d post injection (3 d pIBM) at the time of the second blood feed. *Lp* expression levels were normalized to *Rpl19*. Three biological replicates were analyzed. Means and standard deviation are shown. Unpaired t-test.

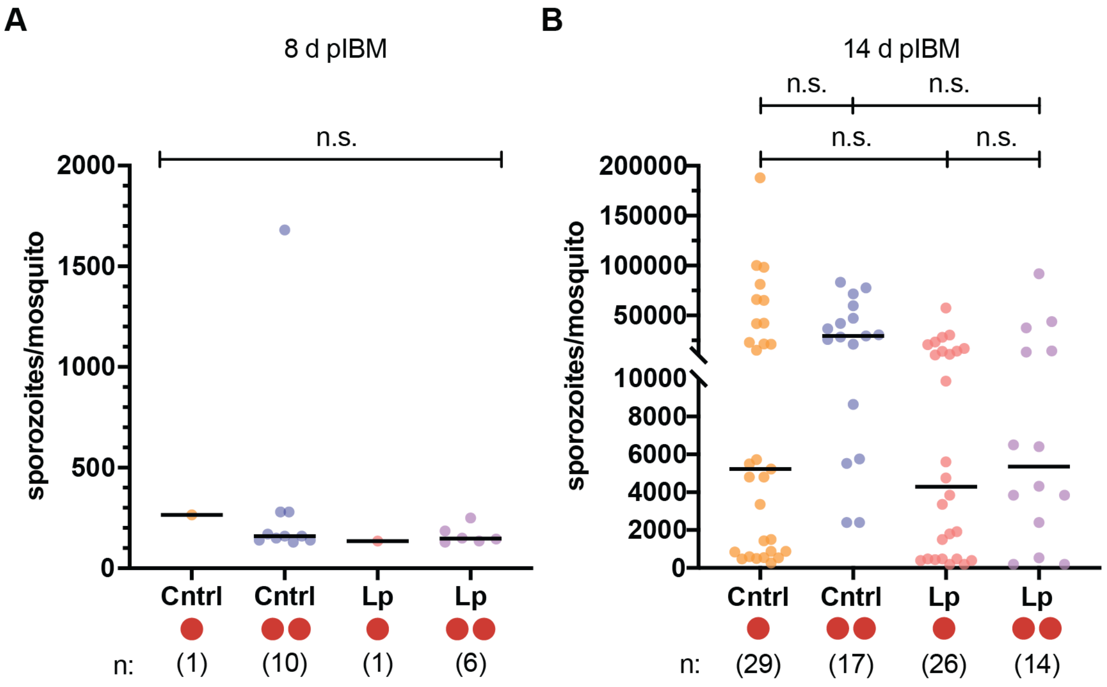


**Fig. S2. Sporozoite intensities in control and Lp-depleted mosquitoes at 8 and 14 d pIBM.** (*A*) Salivary glands of females fed twice (two red circles) show more sporozoites than females fed once (one red circle) at 8 d but low prevalence in singly-fed groups prevents a determination of statistical significance. (*B*) Sporozoite levels in salivary glands at 14 d pIBM are comparable between singly and doubly-fed control and *Lp*-silenced mosquitoes (Linear mixed model; #BF: p=0.0336; ds*RNA*: p=0.0053; FDR-corrected post-hoc Student's t tests shown). Neither the increased infection intensity across the 2BF groups, nor the decreased sporozoite intensity in *Lp*-silenced groups persist after post-hoc testing (Table S1 and S3).

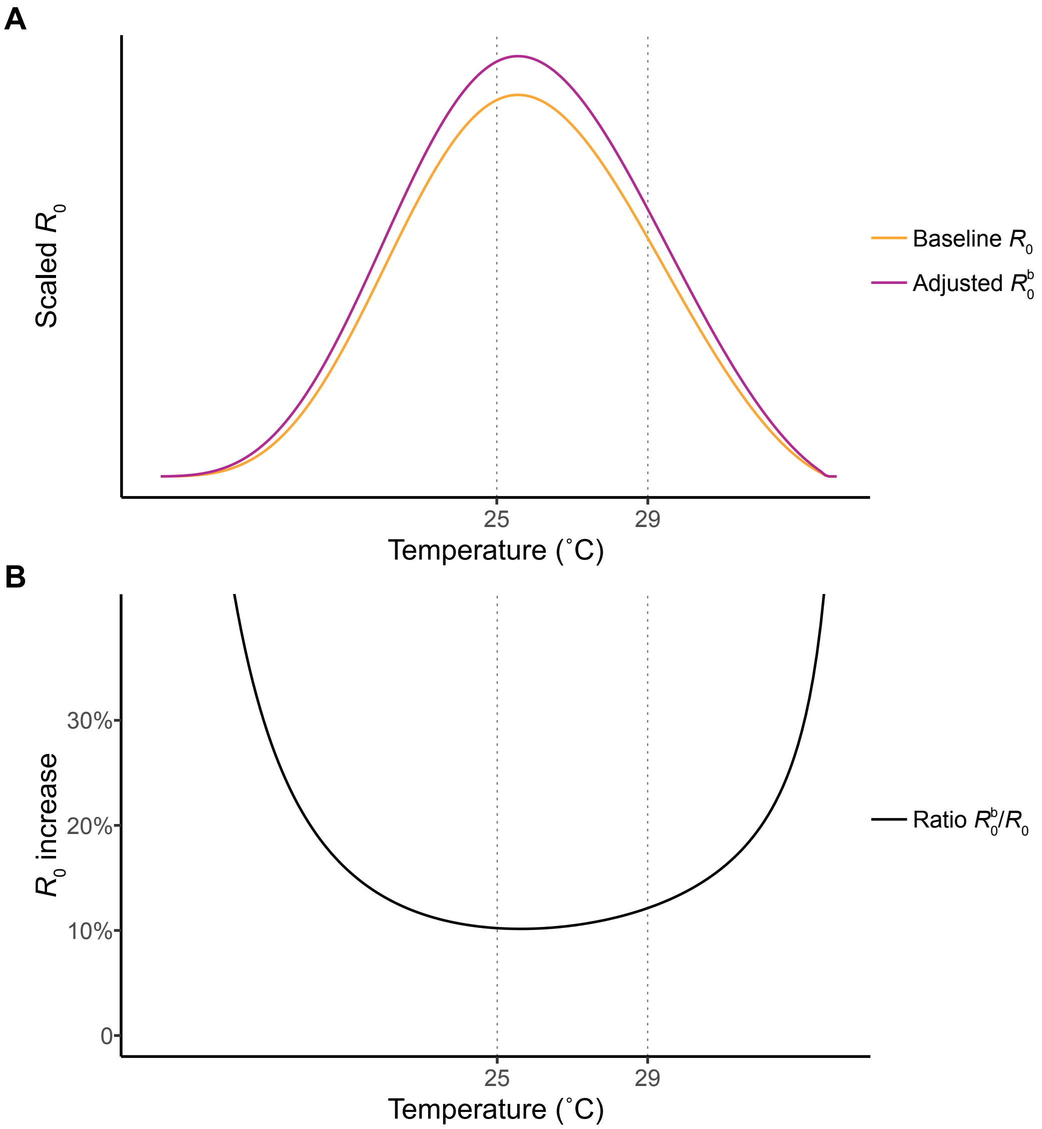


**Fig. S3. The ratio of $R_0$ values is temperature dependent but consistent across temperatures at 27 ± 2 ˚C.** (*A*) Baseline (orange) and Adjusted (red) $R_0$ as temperature varies. No numeric scale is given as raw $R_0$ values depend on parameters, such as population size, that are not temperature dependent and merely cancel out in the $R_0$ ratio. (*B*) The increase in $R_0$ with shortened EIP as a function of temperature. Within the temperature range 27 ± 2 ˚C, the increase in $R_0$ is between 10.1% and 12.1%.

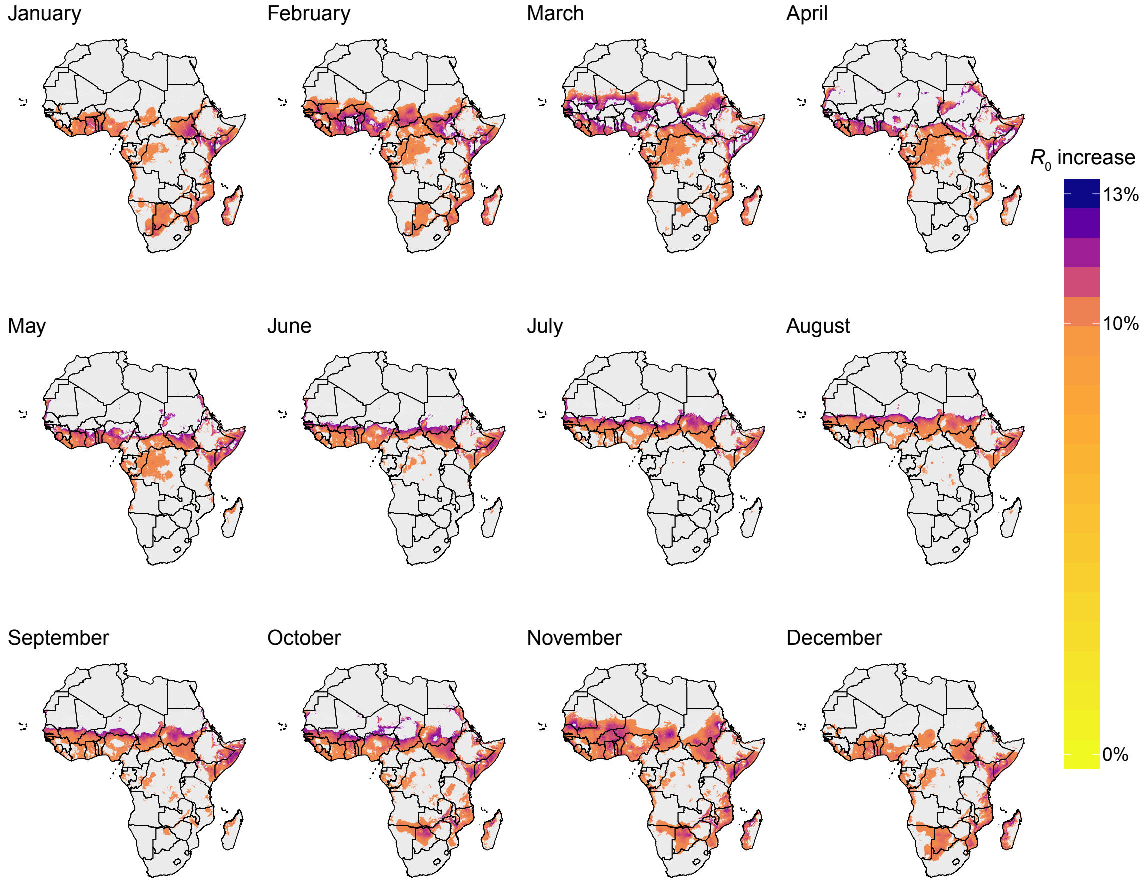


**Fig. S4. Monthly changes in $R_0$ under a multiple blood feeding scenario.** We calculated the monthly changes in $R_0$ by taking the ratio $R_0^b/R_0$ using the mean temperature of each month with a mean temperature 27 ± 2 ˚C for each grid point. The restricted data points shown here are used to create the summary maps in Fig. 4.

**Table S1.** Statistical models. JMP 14 Pro statistical software was used to construct models for data analysis to account for multiple variables in an experiment. Residual Maximum Likelihood (REML) variance components analysis was used by fitting linear mixed models after cube-root transformation to resemble a normal distribution. The number of blood feeds, dsRNA injection group and their interaction were used as fixed effects and replicate was included as a random effect. Effect test outputs are reported here. Multiple comparisons were calculated using 4 pairwise Student's t tests followed by FDR correction (see Table S3). d pIBM = days post infectious blood meal; #BF = number of blood feeds; FDR = false discovery rate.

| Figure | Comparison | Test/Model | Effect Test Outputs |
|---|---|---|---|
| **1B** | 7 d pIBM oocysts,<br>where >0 | Linear Mixed Model<br>followed by 4 post-<br>hoc t-tests, FDR<br>corrected (Table S3) | #BF p=0·5719<br>dsRNA p<0.0001<br>dsRNA x #BF p=0.1733 |
| **1C** | 7 d pIBM<br>mean oocyst size | | #BF p<0·0001<br>dsRNA p=0.1617<br>dsRNA x #BF p=0.3842 |
| **3B** | 10 d pIBM<br>sporozoites,<br>where >0 | | #BF p=0·0001<br>dsRNA p=0.0822<br>dsRNA x #BF p=0.9128 |
| **5A** | 7 d pIBM mean<br>oocyst size | | #BF p<0·0001<br>genotype p<0.0001<br>genotype x #BF p=0.0138 |
| **5B** | 7 d pIBM oocysts,<br>where >0 | | #BF p=0·0409<br>genotype p<0.0001<br>genotype x #BF p=0.7876 |
| **5C** | 10 d pIBM<br>sporozoites,<br>where >0 | | #BF p<0.0001<br>genotype p=0.0056<br>genotype x #BF p=0.2767 |
| **S2A** | 8 d pIBM<br>sporozoites,<br>where >0 | n/a | Insufficient data to run model |
| **S2B** | 14 d pIBM<br>sporozoites,<br>where >0 | Linear Mixed Model<br>followed by 4 post-<br>hoc t-tests, FDR<br>corrected (Table S3) | #BF p=0·0336<br>dsRNA p=0.0053<br>dsRNA x #BF p=0.4303 |

**Table S2.** Statistical testing for infection prevalence. GraphPad Prism 8 was used for logistic regression, Fisher's exact and $\chi^2$ tests.

| Figure | Comparison | Test/Model | Test Outputs |
|---|---|---|---|
| **1B** | 7 d pIBM oocyst prevalence | $\chi^2$ test | $\chi^2$=4.263<br>d.f.=3<br>p>0·05 |
| **3A** | 7 d pIBM sporozoite prevalence | | $\chi^2$=5.697<br>d.f.=3<br>p>0·05 |
| | 8 d pIBM sporozoite prevalence | $\chi^2$ test<br>followed by 4<br>post-hoc $\chi^2$ tests,<br>FDR corrected<br>(Table S4) | $\chi^2$=15.01<br>d.f.=3<br>p=0·0018 |
| | 10 d pIBM sporozoite prevalence | | $\chi^2$=43.33<br>d.f.=3<br>p<0·0001 |
| | 14 d pIBM sporozoite prevalence | $\chi^2$ test | $\chi^2$=2.445<br>d.f.=3<br>p>0·05 |
| **3C** | 7–14d pIBM sporozoite prevalence | Logistic regression<br>Lines of best fit: | Cntrl 1BF: log odds =<br>-8·858+0·8142*d<br>Cntrl 2BF: log odds =<br>-8·646+1·002*d<br>Lp 1BF: log odds =<br>-8.228+0.7125*d<br>Lp 2BF: log odds =<br>-8.762+0.9520*d |
| | | $EIP_{50}$ ± s.e.<br>Z test | Cntrl 1BF: 10.88 ± 0.32 d<br>Cntrl 2BF: 8.63 ± 0.23 d<br>Lp 1BF: 11.55 ± 0.37 d<br>Lp 2BF: 9.20 ± 0.32 d<br>Cntrl 1BF–Cntrl 2BF: z=5.74<br>Cntrl 1BF–Lp 1BF: z=-1.36<br>Cntrl 2BF–Lp 2BF: z=-1.74<br>Lp 1BF–Lp 2BF: z=5.28 |
| **5B** | 7 d pIBM oocyst prevalence | $\chi^2$ test | $\chi^2$=6.488<br>d.f.=3<br>p>0·05 |
| **5C** | 10 d pIBM sporozoite prevalence | $\chi^2$ test,<br>followed by 4<br>post-hoc $\chi^2$ tests,<br>FDR corrected<br>(Table S4) | $\chi^2$=81·43<br>d.f.=3<br>p<0·0001 |
| | 10 d pIBM sporozoite prevalence (pooled by genotype) | Fisher's exact test | p<0·0001 |

**Table S3**. Post-hoc testing for significant differences in oocyst size and oocyst and sporozoite intensity using an FDR of 0.05. See Table S1.

| Fig. 1B | p-value | FDR-adjusted p-value | Significant? |
|---|---|---|---|
| Cntrl 1BF – Cntrl 2BF | $1.80 \times 10^{-1}$ (0.1799) | $2.40 \times 10^{-1}$ (0.2398) | No |
| Cntrl 1BF – Lp 1BF | $7.72 \times 10^{-3}$ (0.0077) | $1.54 \times 10^{-2}$ (0.0154) | Yes |
| Cntrl 2BF – Lp 2BF | $3.21 \times 10^{-5}$ | $1.28 \times 10^{-4}$ (0.0001) | Yes |
| Lp 1BF – Lp 2BF | $5.67 \times 10^{-1}$ (0.5689) | $5.67 \times 10^{-1}$ (0.5689) | No |
| **Fig. 1C** | **p-value** | **FDR-adjusted p-value** | **Significant?** |
| Cntrl 1BF – Cntrl 2BF | $1.15 \times 10^{-24}$ | $2.30 \times 10^{-24}$ | Yes |
| Cntrl 1BF – Lp 1BF | $9.31 \times 10^{-1}$ (0.9309) | $9.31 \times 10^{-1}$ (0.9309) | No |
| Cntrl 2BF – Lp 2BF | $4.36 \times 10^{-2}$ (0.0436) | $5.81 \times 10^{-2}$ (0.0581) | No |
| Lp 1BF – Lp 2BF | $2.8 \times 10^{-32}$ | $1.12 \times 10^{-31}$ | Yes |

| Fig. 3B | p-value | FDR-adjusted p-value | Significant? |
|---|---|---|---|
| Cntrl 1BF – Cntrl 2BF | $3.70 \times 10^{-3}$ (0.0037) | $3.04 \times 10^{-3}$ (0.0148) | Yes |
| Cntrl 1BF – Lp 1BF | $2.66 \times 10^{-1}$ (0.2658) | $2.66 \times 10^{-1}$ (0.2658) | No |
| Cntrl 2BF – Lp 2BF | $1.48 \times 10^{-1}$ (0.1478) | $1.97 \times 10^{-1}$ (0.1971) | No |
| Lp 1BF – Lp 2BF | $6.87 \times 10^{-3}$ (0.0069) | $1.37 \times 10^{-2}$ (0.0137) | Yes |

| Fig. 5A | p-value | FDR-adjusted p-value | Significant? |
|---|---|---|---|
| Cntrl 1BF – Cntrl 2BF | $2.37 \times 10^{-19}$ | $9.48 \times 10^{-19}$ | Yes |
| Cntrl 1BF – Δ*zpg* 1BF | $7.46 \times 10^{-10}$ | $9.95 \times 10^{-10}$ | Yes |
| Cntrl 2BF – Δ*zpg* 2BF | $4.17 \times 10^{-3}$ (0.0042) | $4.17 \times 10^{-3}$ (0.0042) | Yes |
| Δ*zpg* 1BF – Δ*zpg* 2BF | $5.50 \times 10^{-11}$ | $1.10 \times 10^{-10}$ | Yes |
| **Fig. 5B** | **p-value** | **FDR-adjusted p-value** | **Significant?** |
| Cntrl 1BF – Cntrl 2BF | $1.16 \times 10^{-1}$ (0.1156) | $1.54 \times 10^{-1}$ (0.1541) | No |
| Cntrl 1BF – Δ*zpg* 1BF | $3.10 \times 10^{-7}$ | $1.24 \times 10^{-6}$ | Yes |
| Cntrl 2BF – Δ*zpg* 2BF | $1.59 \times 10^{-6}$ | $3.18 \times 10^{-6}$ | Yes |
| Δ*zpg* 1BF – Δ*zpg* 2BF | $5.54 \times 10^{-1}$ (0.5543) | $5.54 \times 10^{-1}$ (0.5543) | No |
| **Fig. 5C** | **p-value** | **FDR-adjusted p-value** | **Significant?** |
| Cntrl 1BF – Cntrl 2BF | $2.73 \times 10^{-5}$ | $5.46 \times 10^{-5}$ | Yes |
| Cntrl 1BF – Δ*zpg* 1BF | $2.09 \times 10^{-2}$ (0.0209) | $2.79 \times 10^{-2}$ (0.0279) | Yes |
| Cntrl 2BF – Δ*zpg* 2BF | $1.13 \times 10^{-1}$ (0.1126) | $1.13 \times 10^{-1}$ (0.1126) | No |
| Δ*zpg* 1BF – Δ*zpg* 2BF | $3.14 \times 10^{-6}$ | $1.26 \times 10^{-5}$ | Yes |

| Fig. S2B | p-value | FDR-adjusted p-value | Significant? |
|---|---|---|---|
| Cntrl 1BF – Cntrl 2BF | $3.10 \times 10^{-2}$ (0.0310) | $6.20 \times 10^{-2}$ (0.0620) | No |
| Cntrl 1BF – Lp 1BF | $8.74 \times 10^{-2}$ (0.0874) | $1.16 \times 10^{-1}$ (0.1165) | No |
| Cntrl 2BF – Lp 2BF | $2.43 \times 10^{-2}$ (0.0243) | $9.72 \times 10^{-2}$ (0.0972) | No |
| Lp 1BF – Lp 2BF | $3.51 \times 10^{-1}$ (0.3510) | $3.51 \times 10^{-1}$ (0.3510) | No |

**Table S4.** Post-hoc testing for significant differences in infection prevalence. See Table S2.

| **Fig. 3A – 8 d pIBM** | **p-value** | **FDR-adjusted p-value** | **Significant?** |
|---|---|---|---|
| Cntrl 1BF – Cntrl 2BF | $2.56 \times 10^{-3}$ (0.0026) | $1.04 \times 10^{-2}$ (0.0104) | Yes |
| Cntrl 1BF – Lp 1BF | 1.00 | 1.00 | No |
| Cntrl 2BF – Lp 2BF | $3.85 \times 10^{-1}$ (0.3848) | $5.13 \times 10^{-1}$ (0.5131) | No |
| Lp 1BF – Lp 2BF | $5.17 \times 10^{-3}$ (0.0517) | $1.03 \times 10^{-1}$ (0.1034) | No |
| **Fig. 3A – 10 d pIBM** | **p-value** | **FDR-adjusted p-value** | **Significant?** |
| Cntrl 1BF – Cntrl 2BF | $2.19 \times 10^{-6}$ | $8.76 \times 10^{-6}$ | Yes |
| Cntrl 1BF – Lp 1BF | $3.29 \times 10^{-1}$ (0.3289) | $3.29 \times 10^{-1}$ (0.3289) | No |
| Cntrl 2BF – Lp 2BF | $1.69 \times 10^{-1}$ (0.1687) | $2.23 \times 10^{-1}$ (0.2253) | No |
| Lp 1BF – Lp 2BF | $2.10 \times 10^{-5}$ | $4.20 \times 10^{-5}$ | Yes |
| **Fig. 3C – $\Delta EIP_{50}$** | **p-value** | **FDR-adjusted p-value** | **Significant?** |
| Cntrl 1BF – Cntrl 2BF | $9.47 \times 10^{-9}$ | $3.79 \times 10^{-8}$ | Yes |
| Cntrl 1BF – Lp 1BF | $1.74 \times 10^{-1}$ (0.1740) | $1.74 \times 10^{-1}$ (0.1740) | No |
| Cntrl 2BF – Lp 2BF | $8.19 \times 10^{-2}$ (0.0819) | $1.09 \times 10^{-1}$ (0.1092) | No |
| Lp 1BF – Lp 2BF | $1.29 \times 10^{-7}$ | $2.58 \times 10^{-7}$ | Yes |

| **Fig. 5C** | **p-value** | **FDR-adjusted p-value** | **Significant?** |
|---|---|---|---|
| Cntrl 1BF – Cntrl 2BF | $1.42 \times 10^{-11}$ | $4.26 \times 10^{-11}$ | Yes |
| Cntrl 1BF – Δ*zpg* 1BF | $5.74 \times 10^{-9}$ | $1.15 \times 10^{-8}$ | Yes |
| Cntrl 2BF – Δ*zpg* 2BF | $1.52 \times 10^{-1}$ (0.1517) | $1.82 \times 10^{-1}$ (0.1820) | No |
| Δ*zpg* 1BF – Δ*zpg* 2BF | $2.76 \times 10^{-3}$ (0.0028) | $4.14 \times 10^{-3}$ (0.0041) | Yes |

**Table S5.** Temperature dependent traits fitted with Briere function. All parameters from Mordecai *et al.* (1). See references within.

| Trait | $c$ | $T_0$ (°C) | $T_m$ (°C) |
|---|---|---|---|
| *a* | 0.000203 | 11.7 | 42.3 |
| *PDR* | 0.000111 | 14.7 | 34.4 |
| *MDR* | 0.000111 | 14.7 | 34 |

**Table S6.** Temperature dependent traits fitted with quadratic function. All parameters from Mordecai *et al.* (1). See references within.

| **Trait** | ***q*** | ***r*** | ***s*** |
|---|---|---|---|
| *bc* | -0.54 | 25.2 | -206 |
| *p* | -0.000828 | 0.0367 | 0.522 |
| $p_{EA}$ | -0.00924 | 0.453 | -4.77 |
| *EFD* | -0.153 | 8.61 | -97.7 |
